# SIC50: Determination of IC50 by an optimized Sobel operator and a vision transformer

**DOI:** 10.1101/2022.08.12.503661

**Authors:** Yongheng Wang, Weidi Zhang, Hoyin Yip, Chuyuan Qu, Hongru Hu, Xiaotie Chen, Teresa Lee, Xi Yang, Bingjun Yang, Priyadarsini Kumar, Su Yeon Lee, Javier J. Casimiro, Jiawei Zhang, Kit S. Lam, Aijun Wang

## Abstract

As a measure of cytotoxic potency, half-maximal inhibitory concentration (IC50) is the concentration at which a drug exerts half of its maximal inhibitory effect against target cells. It can be determined by various methods that require applying additional reagents or lysing the cells. Here, we describe a label-free Sobel-edge-based method, which we name SIC50, for the evaluation of IC50. SIC50 classifies pre-processed phase-contrast images with a state-of-art vision transformer and allows for the continuous assessment of IC50 in a faster and more cost-efficient manner. We have validated this method using four drugs and 1536-well plates, as well as built a first-of-its-kind web application. We anticipate this method will assist in the high-throughput screening of chemical libraries (e.g., small molecule drugs, siRNA, and microRNA and drug discovery.

---

One in three people will be diagnosed with a type of cancer in their lifetime^1^. In 2019, $141.3 billion was spent on oncology medicines globally. This number was projected to reach $394.2 billion by 2027^2^. The development of cancer therapeutics is expensive, costing approximately $648 million per drug in 2017^3^, partially due to the necessary screening of tens of thousands of chemicals, which can be prohibitively costly. Thus far, many clinical trials have failed because they were not able to start with the best possible molecule. Therefore, a more affordable method for the efficient evaluation of drug efficacy and the screening of cancer therapeutics is urgently needed to advance the field of cancer treatment.

Current methods require chemicals and several steps to evaluate cell viability and drug potency. For example, developed in 1983, the 3-(4,5-dimethylthiazol-2-yl)-2,5-diphenyl-2H-tetrazolium bromide (MTT) assay has been widely adopted to determine the efficacy of drugs (Fig. 1a)^4^. The positively-charged, yellow MTT molecules penetrate viable cells and become reduced to purple formazan crystals by mitochondrial dehydrogenase^5–6^. The crystal is then dissolved in dimethyl sulfoxide (DMSO, 100 μL/well) or sodium dodecyl sulfate (SDS 10% w/v in 0.01M hydrochloric acid, 100 μL/well) to become a colored solution with an absorbance maximum near 570 nm, which is quantified by a spectrophotometer. The absorbance is proportional to the number of live cells. The darker the solution, the greater the number of viable cells. Thus, this colorimetric assay measures metabolic activity as an indicator of cell viability^7^. Cell counting kit-9 (CCK-8) is another commonly used assay. It is based on the reduction of water-soluble tetrazolium 8 (WST-8) to 1-methoxy phenazinium methylsulfate (PMS) by NAD- or NADP-dependent dehydrogenases in a cell (Fig. 1b). Compared to MTT, CCK-8 is more sensitive and does not require dissolving solutions^8^. In an adenosine triphosphate (ATP) assay, cells are lysed to release the ATP, which activates luciferin and yields a luciferyl-adenylate and pyrophosphate. The luciferyl-adenylate reacts with oxygen to generate carbon dioxide and oxyluciferin in an electronically excited state, which releases bioluminescence (550-570nm) when returned to the ground state. The luminescent signal is proportional to the ATP levels and the number of viable cells (Fig. 1c). Lastly, the IC50 of a drug can also be determined by counting the nuclei stained with Hoechst dye. However, we observed that the staining inhibited the growth of cells (Fig. 1d).

**Fig. 1.**
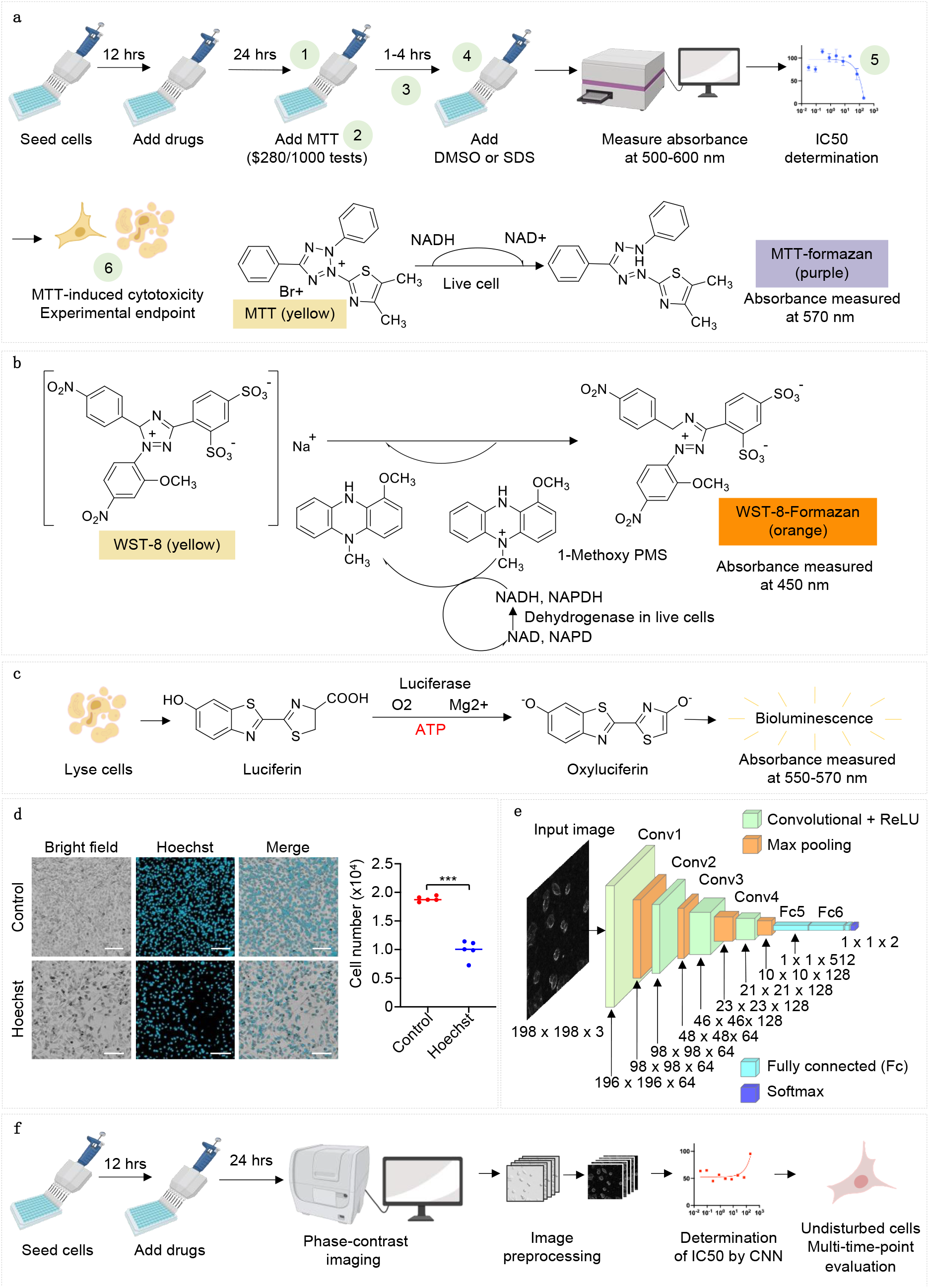
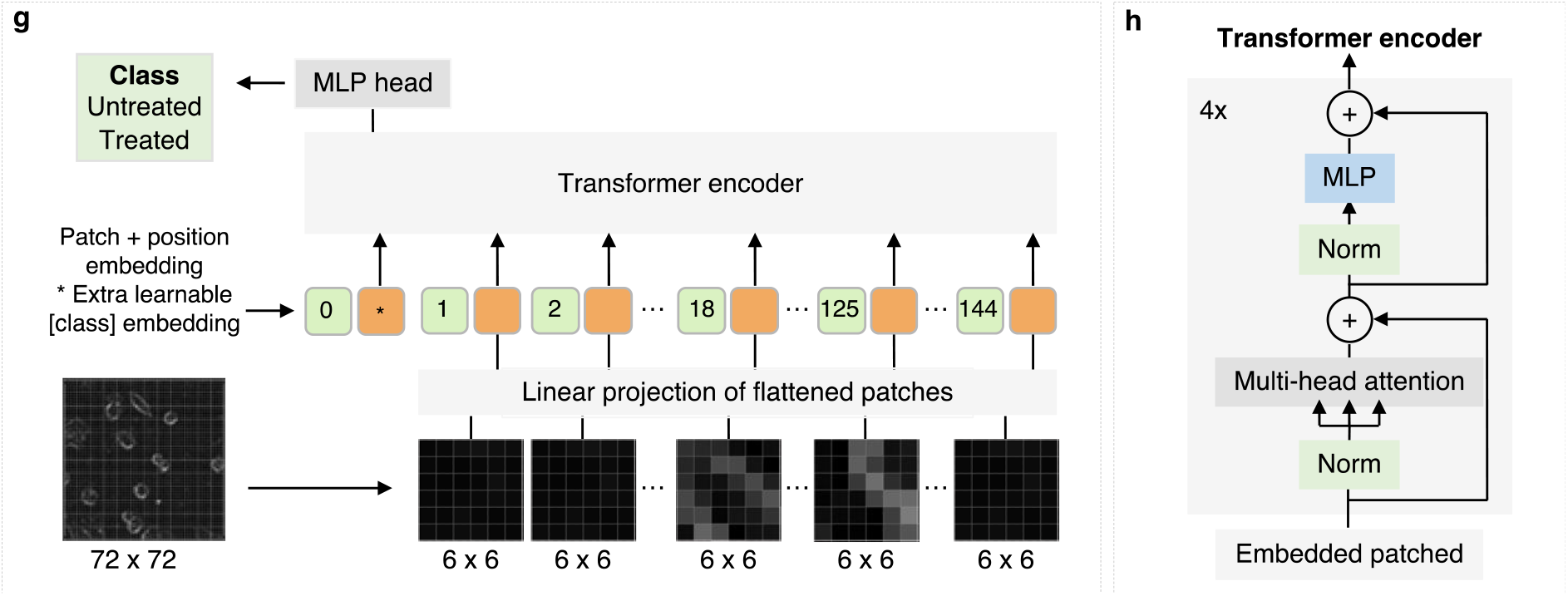
Comparison of SIC50 and commonly used cell viability assays. **a,** The workflow for an MTT assay. MTT (0.2-0.5 mg/mL) is reduced to its insoluble formazan by nicotinamide adenine dinucleotide phosphate (NADPH)-dependent oxidoreductase in metabolically active cells. The absorbance of MTT-formazan is measured at 570nm. **b,** Cell counting kit-8 (CCK-8) assays rely on the reduction of water-soluble tetrazolium 8 (WST-8) to 1-methoxy phenazinium methylsulfate (PMS). **c,** In ATP assays, cells are lysed to release ATP. ATP helps activate luciferin and generate a luciferyl-adenylate and pyrophosphate. Luciferyl-adenylate will be oxidized and yield electronically excited oxyluciferin, which will return to the ground state and release yellow luminescent light. **d,** Hoechst inhibits the growth of cells. Images are obtained from B16-F10 cells. In the control group, the cells were stained with 5 μg/mL Hoechst 33342 for 20 minutes. In the Hoechst group, the cells were incubated with Hoechst for 24 hours. Scale bars are 40 μm. Statistical analyses of the cell numbers from the control group and the Hoechst group suggest that the two groups are significantly different. n=5, Mann-Whitney test, p=0.008. **e,** The architecture of the Conv2D used in this study. **f,** The procedures of SIC50. SIC50 determines the IC50 values of drugs by analyzing phase-contrast images modified with a Sobel operator. It avoids operational errors, saves time and reagent cost, will not be interfered by chemicals with absorbance between 450 nm and 600 nm, and allows for multi-time-point assessments of IC50. **g-h**, The overview of a vision transformer. Images are split into patches, position-embedded, and fed into a vision transformer encoder. An extra learnable embedding is added for classification.

Deep learning has been used to analyze histological images^9–10^, study the differentiation of induced pluripotent stem cells (iPSCs)^11–15^, and perform binary classifications (live or dead^16^, drug-treated or -untreated^17^) on cancer cells. As a self-attention-based deep neural network, a transformer was developed in 2017^18^ for tasks in the field of natural language processing and employed in computer vision applications since 2021^19^, thanks to its strong representation capabilities and less need for vision-specific inductive bias.

In this study, a vision transformer and convolutional neural networks (Conv2D) were built to classify preprocessed phase-contrast images and predict the IC50 of drugs (Fig. 1e-h). Compared to the widely-used MTT assay, our method has the following advantages: 1) avoids operational errors and saves time associated with adding the reagents; 2) reduces costs associated with the labeling reagent (i. e. MTT), balancing buffer, filters, and dissolving solutions; 3) does not require incubation time; 4) can be used to screen a broader range of chemicals because compounds with absorbance from 450nm to 600nm or with antioxidant properties will interfere with the MTT or CCK-8 absorbance measurement^20–21^; 5) the cells do not need to be in the log phase using our method. In MTT assays, however, cells need to be in the log phase to ensure the linearity between absorbances and cell numbers; 6) our method is not cytotoxic, permitting multi-time-point measurements (Fig. 1f).

Binary classification is conducted using untreated cells as the control. The classification accuracies are low before preprocessing the images (Fig. 2a) or modifying the images with a high-pass algorithm (Fig. 2b). The accuracy is significantly improved when the images are processed with a Sobel operator (Fig. 2c) and an optimized Sobel (OSobel) operator (Fig. 2d), likely due to the operator reducing the grayscale values of the background and increasing the signal-to-noise ratio (Fig. 2a and 2d). Regardless of the preprocessing methods, the accuracies are equally poor at low concentrations for treated cells (Fig. 2e-f). However, when the cells are treated with drugs (e.g., cephalotazine and fasudil) at high concentrations, the classification accuracies using images preprocessed with an OSobel algorithm are higher than that of Gaussian high pass and unprocessed images (Fig. 2e-f).

**Fig. 2.**
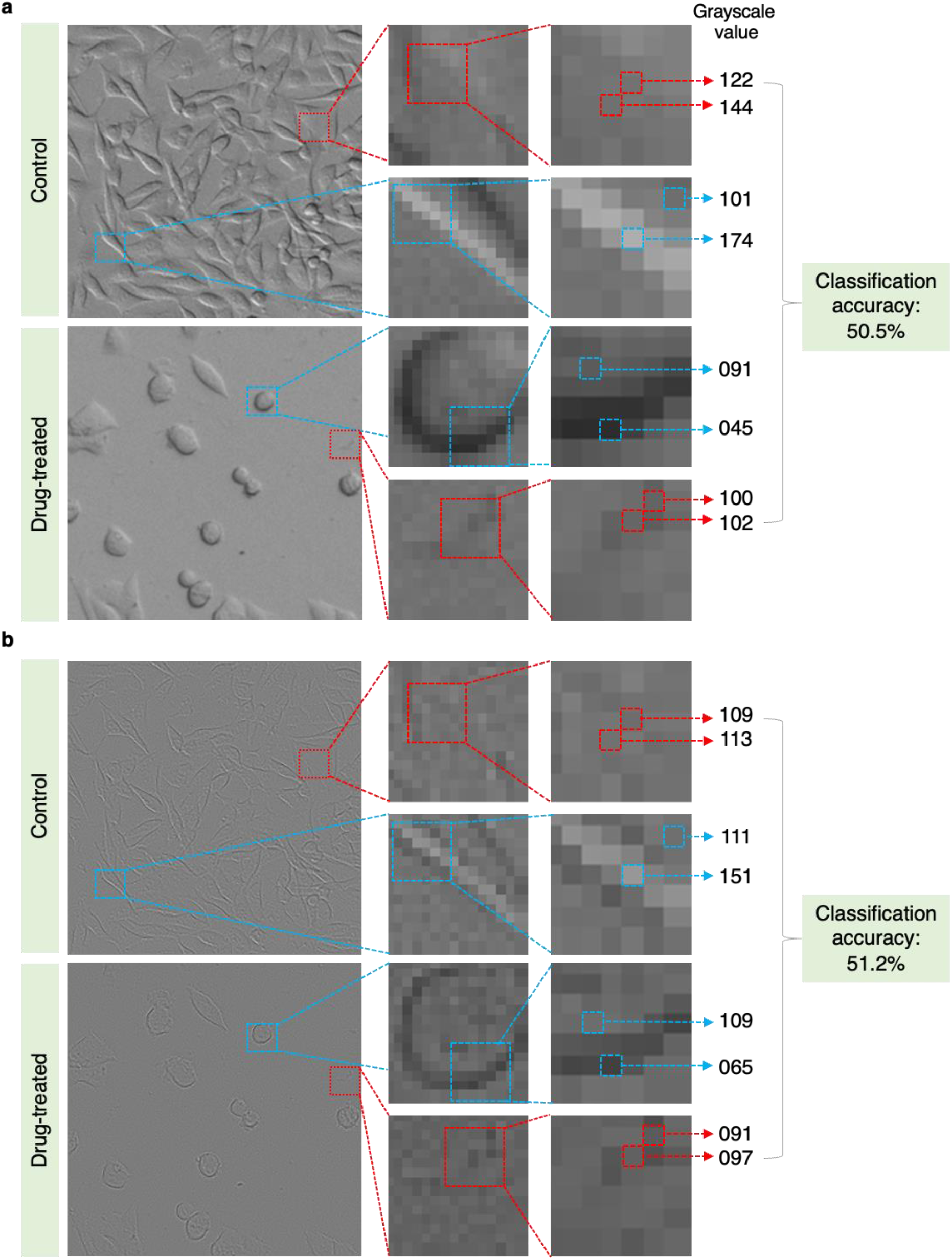
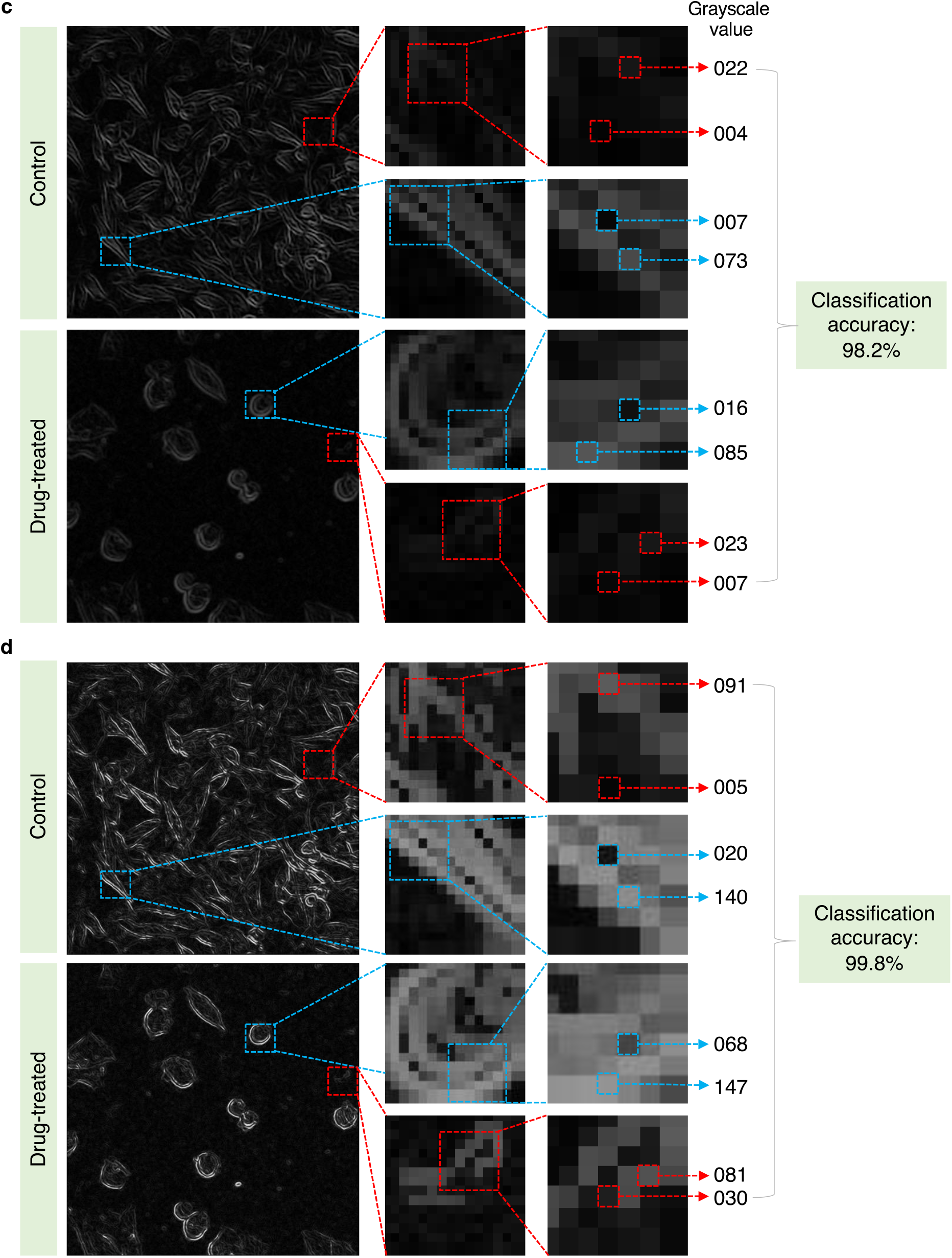
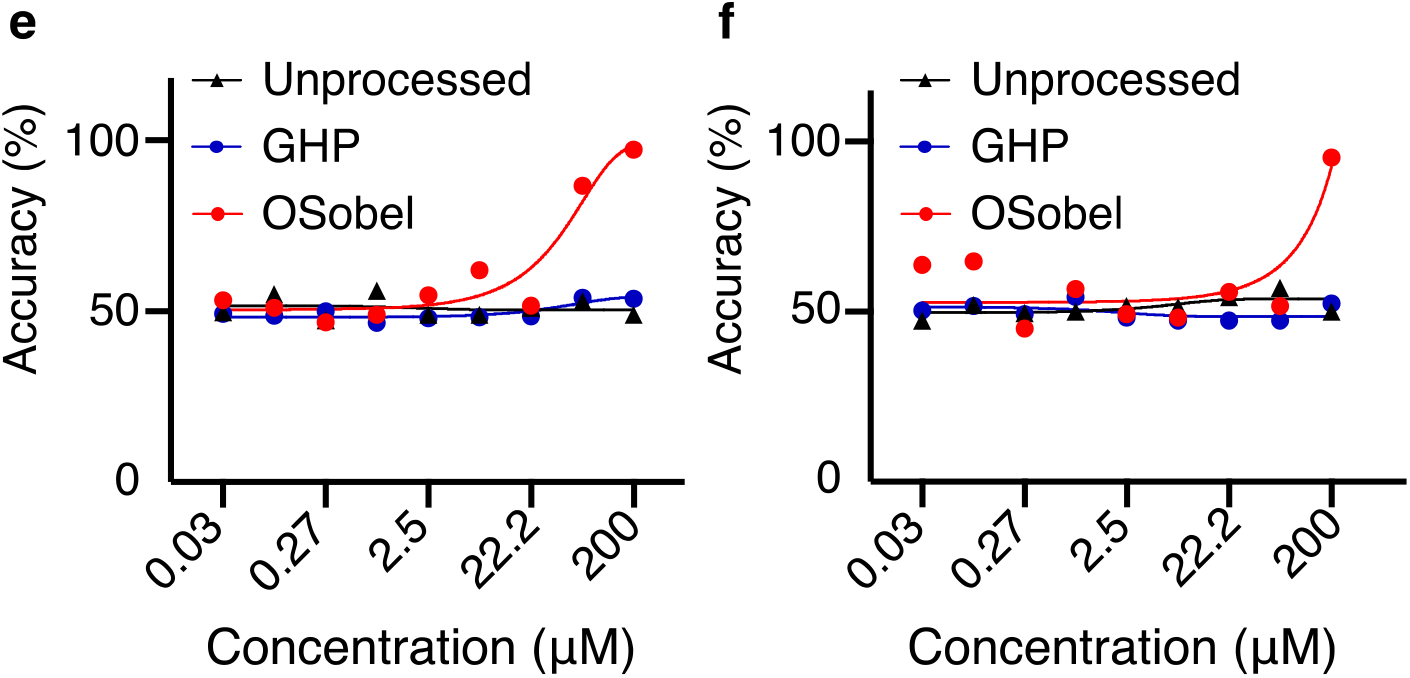
Optimized Sobel operator improves classification accuracy and outperforms other methods. **a,** The B16-F10 cells were treated with 200 μM paclitaxel for 24 hours. Conv2D can not accurately classify untreated and treated cells before preprocessing images (accuracy=50.5%). The grayscale values of the background are large and relatively close to that of cells. **b,** Gaussian high-pass (GHP) does not significantly improve the prediction accuracy (accuracy=51.2%). The difference between background and signal is small. **c,** Sobel filter reduces the grayscale values of the background, increases the signal-to-noise ratio, and improves prediction accuracy to 98.2%. **d,** An optimized Sobel (OSobel) operator further improved the prediction accuracy to 99.8%. R, red. G, green. B, blue. **e,** The classification accuracies of Conv2D using cells treated with cephalotaxine at various concentrations. The red curve is the same as the one in Fig. 3c. **f,** The classification accuracies of Conv2D using cells treated with different concentrations of fasudil. The red curve is the same as the one in Fig. 3d.

We observe that the accuracy of the binary classification can be used to predict the IC50 of a drug (Fig. 3a). As a reference, the cell viability and IC50 of drugs are first determined using Hoechst staining and represented as blue dots. Binary classifications are performed using untreated cells and the cells treated with drugs at various concentrations. The classification accuracies at different concentrations are labeled as red squares. The accuracies of binary classification are close to 50% when the cells are treated with drugs at low concentrations, suggesting that the drugs do not result in detectable changes in cellular morphology. Drawing lines connecting every pair of adjacent red squares, the line with the highest slope contains values close to the IC50s of drugs (Fig. 3b-f). Although the average value of these two adjacent concentrations (SIC50) can be different from the IC50s estimated using Hoechst staining by 2-fold (HIC50, Table 1), mapping the IC50 from a large range of concentrations (0.03 μM to 200 μM, 6666-fold) to a 2-fold variation can be helpful and sufficient for many purposes. In addition, 1.5-fold or 1.25-fold dilutions could be adopted to further improve the prediction accuracy of SIC50. Although the classification accuracies using a vision transformer are close to that of Conv2D (Fig. 3b–3e), we expect the vision transformer would perform better when pre-trained at a large scale and transferred to tasks with fewer images^19^.

**Fig. 3.**
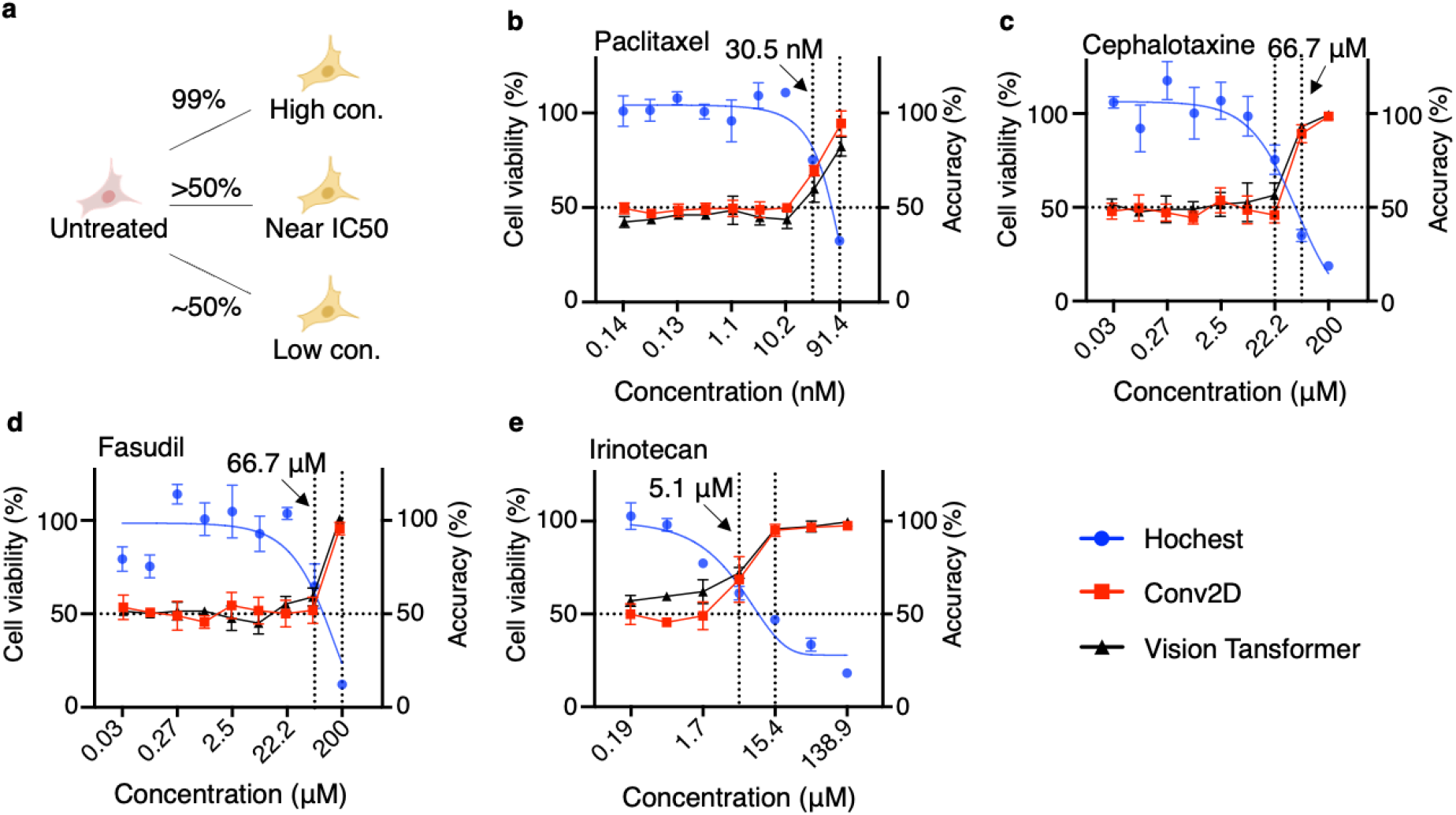
Determination of IC50s of drugs by SIC50. **a,** Binary classifications are performed using the untreated cells as a control. The classification accuracy suddenly increases when the cells are treated with a drug near its IC50. **b-e,** The cell viability (blue curves, left y-axis) and classification accuracies (red curves, right y-axis) in cells treated with drugs at various concentrations respectively, including paclitaxel (b) at 91.4 nM, 30.5 nM, 10.2 nM, 3.4 nM, 1.1 nM, 0.38 nM, 0.13 nM, 0.04 nM and 0.014 nM, and cephalotaxine (c) and fasudil (d) at 200 μM, 66.7 μM, 22.2 μM, 7.4 μM, 2.5 μM, 0.82 μM, 0.27 μM, 0.09 μM, and 0.03 μM and irinotecan (e) at 138.9 μM, 46.3 μM, 15.4 μM, 5.1 μM, 1.7 μM, 0.57 μM, and 0.19 μM.

**Table 1.** IC50 values determined by Hoechst staining and Conv2 and a vision transformer

|  |  |  |  |
| --- | --- | --- | --- |
| IC50s | Hoechst (HIC50) | SIC50 | Difference |
| Paclitaxel (nM) | 31.50 | 60.91 | 1.93 |
| Cephalotaxine (μM) | 32.94 | 44.45 | 1.35 |
| Fasudil (μM) | 69.27 | 133.35 | 1.93 |
| Irinotecan (μM) | 6.03 | 10.25 | 1.70 |

The SIC50 method is validated using four drugs whose mechanisms of action are different. Paclitaxel stabilizes microtubules, increases microtubule polymerization, decreases microtubule depolymerization, prevents mitosis, and blocks cell cycle progression^22^. Cephalotaxin inhibits the growth of cancer cells by activating the mitochondrial apoptosis pathway^23^. Fasudil is a calcium channel blocker and inhibits the Rho-kinase signaling pathway^24–25^. Irinotecan is a prodrug of 7-ethyl-10-hydroxycamptothecin (SN-38), which forms complexes with topoisomerase 1B and DNA, and causes DNA misalignment and cell death^26^.

Although deep learning has been employed for binary classification of cancer cells (live or dead^16^, drug-treated or -untreated^17^), the cells were treated with the drug at only one concentration^16^. The classification accuracies did not change significantly when the cells were treated for 48 hours probably due to differences in preprocessing methods^17^. Thus, the accuracies were not used for the assessment of IC50.

Validated in four drugs, we anticipate our method will empower drug discovery and research in pharmacology by facilitating the high-throughput screening of chemical libraries using 1536-well plates and Cytation 5, helping evaluate the potency of other small molecule drugs, siRNA, and microRNA, and assessing the cytotoxicity of delivery vehicles for drugs and genes such as lipid nanoparticles^27^, polymers, and adeno-associated viruses^28–31^. In addition, we envision our method can be modified to facilitate biomedical research related to changes in cellular morphology, e.g., cancer cell metastasis, stem cell differentiation, neural plasticity, and so on.

## Materials and Method

The following materials were procured from different sources: paclitaxel (LC laboratories, Cat. P-9600, CAS No. 33069-62-4), cephalotaxine (Toronto Research Chemicals, Cat. C261050, CAS No. 24316-19-6), irinotecan (APExBIO, Cat. B2293, CAS No. 136572-09-3), fasudil (Toronto Research Chemicals, Cat. R1036), cantharidin (Sigma Aldrich, Cat. C7632, CAS No. 56-25-7), B16-F10 cells (ATCC, Cat. CRL-6475), Dulbeccos modified eagle medium (DMEM, Fisher Scientific, Cat. 11-995-073), fetal bovine serum (FBS, Fisher Scientific, Cat.10-437-028), penicillin-streptomycin (PS, Thermo Fisher Scientific, Cat. 15070063), 96-well plates (Corning, product No.3595 and Fisher Scientific, Cat. No. FB012931), 1536-well plates (Corning 3712), 384-well plates (Corning 3833), Hoechst 33342 (Thermo Fisher Scientific, Cat. No. H3570), and an automated cell imaging system (Molecular Devices, ImageXpress Pico).

### Cell culture

B16-F10 cells were seeded in plates (96-well plates at the density of 6000 cells/well, 384 well plates at the density of 1800 cells/well, and 1536-well plates at the density of 240 cells/well) in complete growth media (DMEM, 10% FBS, and 1% penicillin/streptomycin) and cultured for 12 hours at 37 °C incubators containing 5% CO_2_. Drugs were subsequently added at various concentrations and incubated for additional 24 hours.

### Determination of IC50 by Hoechst staining

The cells are incubated with culture media containing 1 μg/ml Hoechst 33342 for 10 minutes. The IC50 is calculated after counting the number of nuclei with CellProfiller and curve fitting with GraphPad Prism 9. Approximately, the IC50 equals the x-axis value of the intersecting point of a blue curve in Fig. 3 and the horizontal dashed line crossing the y-axis at 50%.

### Image collection and preprocessing

Images are collected by Pico (4X objective, 29.8% of well), exported as 16-bit TIFF, batched converted into 8-bit TIFF, then into 8-bit PNG by ImageJ, and processed using different algorithms (Code posted on GitHub). Afterward, each image is split into 100 smaller images by PhotoScape X, followed by data augmentation to generate more images for training.

### Architecture of Conv2D

The Conv2D is constructed using the TensorFlow framework. The network consists of 4 convolutional layers with the rectified linear unit (ReLU). The first two convolutional layers have 64 kernels, while the third and the fourth convolutional layers have 128 kernels. At each convolutional layer, the convolution is performed by sliding the filter over the input data to extract the hidden features from the data. Each convolutional layer is followed by a 2*2 max pooling layer to reduce the feature dimension by keeping only the most relevant features. The output from the last pooling layer is then flattened and presented to a dropout layer with a rate of 0.5 to avoid overfitting. The last 2 layers in the Conv2D are fully connected layers, with 512 and 2 neurons respectively, followed by a to obtain the final classification result. The parameters of the network are trained over 50 epochs randomly using the training data (90% for training and 10% for test). The accuracy of the classifier is then computed by evaluating the fitted model using the test data.

### Architecture of a vision transformer

Images are resized to 72 x 72 and split into 6 x 6 patches to build a transformer encoder together with position embeddings and a learnable “classification token”. Layer normalization is implemented before every block of the transformer encoder containing multiheaded self-attention and multilayer perceptrons (MLPs). Each MLP consists of two layers with Gaussian error linear unit (GELU) non-linearity. We use a learning rate of 0.001 and a weight decay of 0.0001, a batch size of 200, 100 epochs, 4 transformer layers, a dropout rate of 0.1, and MLP head units of [2048, 1024].

### Code, data, and web application

The code and images used in this study are uploaded to GitHub (https://github.com/BioChemML/SIC50) and Google Drive (https://drive.google.com/drive/folders/1V_krsQRlg0vAJPfd88dyuR_nz3sLf2Fy?usp=sharing), respectively. The link to the web application is http://biochemml.com/image/

## Author contribution

Y.W. trained the models, built the web application, and wrote the first draft. W. Z. facilitated data collection and analyses. H. Y. C. Q., H. H., B. Y., T. L., X. C., P. K., S., J. J. C, Y., J. Z., K. S. L., and A. W. discussed the project and reviewed the manuscript.

## Acknowledgments

This work was supported by the Science Translation and Innovative Research (STAIR) grant offered by the University of California Davis Venture Catalyst. Fig. 1a, 1c, and 1 f, and Fig. 3a were created with BioRender.com. Fig. 1a, 1b, and 1c were produced using ChemDraw. Fig. 1e was generated on app.diagrams.net. We thank Alexandra M. Iavorovschi for proofreading the manuscript.

